# Single-cell transcriptomics showed that maternal PCB exposure dysregulated ER stress-mediated cell type-specific responses in the liver of female offspring

**DOI:** 10.1101/2025.06.04.657944

**Authors:** Joe Lim, Youjun Suh, Xueshu Li, Rebecca Wilson, Hans-Joachim Lehmler, Pamela Lein, Julia Yue Cui

**Author notes:** Address correspondence to: Julia Yue Cui, PhD, DABT Professor Department of Environmental and Occupational Health Sciences University of Washington.

## Abstract

Polychlorinated biphenyls (PCBs) are persistent environmental toxicants that bioaccumulate in the food chain and readily cross the placenta, raising concerns for developmental toxicity. While PCB exposure has been associated with metabolic and neurodevelopmental disorders, its cell type-specific effects on liver development remain poorly understood. This study aimed to investigate how maternal exposure to an environmentally relevant Fox River PCB mixture affects liver development in female offspring at single-cell resolution. We hypothesized that early-life PCB exposure disrupts hepatic metabolic and immune function in a cell type-specific manner. Using single-cell RNA sequencing (scRNA-seq) on liver tissue from postnatal day 28 female mice perinatally exposed to PCBs, we identified major hepatic and immune cell populations and assessed cell-specific transcriptional responses. PCB exposure significantly altered the proportions of endothelial cells and Kupffer cells and reduced neutrophil abundance. Transcriptomic analysis revealed that PCBs dysregulated key pathways in hepatocytes and non-parenchymal cells, including ER stress responses, drug metabolism, and glucose/insulin signaling. Notably, hepatocytes exhibited upregulation of phase-I drug-metabolizing enzymes and uptake transporters, but downregulation of phase-II enzymes and efflux transporters. Kupffer cells and endothelial cells had altered immune and metabolic gene expression, and intercellular communication analysis predicted disrupted fibronectin, collagen, and chemokine signaling due to PCB exposure. RT-qPCR validation confirmed increased hepatic ER stress marker expression. Together these findings demonstrate that perinatal PCB exposure induces persistent, cell type-specific transcriptomic reprogramming in the liver, impairing metabolic and immune functions. This study highlights the utility of single-cell transcriptomics for revealing toxicant effects with cellular precision during critical windows of development.

## INTRODUCTION

Polychlorinated biphenyls (PCBs) are persistent organic compounds that were widely used as electrical coolants and insulators, hydraulic fluids, and plasticizers (Erickson and Kaley, 2011). Although PCBs have been banned from commercial production in the United States in the 1970s (Ross, 2004) and global production in the 2000s (Klocke and Lein, 2020), they are still present in the environment and bioaccumulate in food chains (Melymuk *et al*., 2022) and (Zhu *et al*., 2022), which raises safety concerns for continuous exposure. PCBs were detected in various food items, such as meat, milk, fish, and eggs (Shen *et al*., 2017; Saktrakulkla *et al*., 2020; Stadion *et al*., 2024). In addition, PCBs can cross the placental barrier and are excreted through breast milk (Jacobson *et al*., 1984; Winneke *et al*., 2002; Wang *et al*., 2004). Therefore, early life is a sensitive window of PCB exposure.

Exposure to PCBs is linked to immune system dysfunction (Montano *et al*., 2022), reproductive toxicity (Casas *et al*., 2015; Montano *et al*., 2022), cardiovascular diseases (Fiolet *et al*., 2021), and neurodevelopmental toxicity (Bullert *et al*., 2021). Epidemiology studies showed that developmental exposure to PCBs is linked to cognitive development and attention issues (Klocke and Lein, 2020; Balalian *et al*., 2024). In rodent models, perinatal exposure to PCBs resulted in reduced vocalizations, increased repetitive behavior, and social deficits (Sethi *et al*., 2021), gut barrier defects and increased inflammation (Rude *et al*., 2019), as well as endocrine disruption including dysregulation of estrogen receptor alpha signaling (Dickerson *et al*., 2011). Developmental exposure to the Fox River PCB mixture, an environmentally-relevant PCB mixture (Kostyniak *et al*., 2005), impairs social communication related to neurodevelopmental disruption in mice (Wilson *et al*., 2024). Exposure to the Fox River PCB mixture during pregnancy altered metabolites involved in glucuronidation and heme and amino acid metabolism in serum, suggesting potential contribution to developmental neurotoxic effects of the pups (Li *et al*., 2024). The liver is a major organ for xenobiotic biotransformation and nutrient homeostasis (McLauchlan, 1997) and is a target organ for PCB exposure. Previous work showed that PCB exposure is positively associated with prevalence of metabolic disorders and liver diseases in human populations (Clair *et al*., 2018; Cave *et al*., 2022). In laboratory studies, PCB exposure resulted in lipid accumulation and steatohepatitis signatures in rats (Vieira Silva *et al*., 2022). PCB exposure led to hepatic fibrosis, inflammation, and metabolic dysfunction, which were exacerbated from diet-induced liver injury (Wahlang, Barney, *et al*., 2017; Wahlang, Perkins, *et al*., 2017; Deng *et al*., 2019). We previously showed that acute exposure to the Fox River PCB mixture altered bile acid homeostasis within the gut-liver axis (Cheng *et al*., 2018), and that the gut microbiome is necessary for PCB-induced changes in the hepatic transcriptome including dysregulation of xenobiotic and steroid metabolism in adult mice (Lim *et al*., 2020). However, the effects of PCB exposure on liver development during early life—particularly its impact on metabolic and immune regulatory pathways—remain poorly understood. In particular, how PCBs influence liver maturation at **single-cell resolution** and with **cell-type-specific precision** during critical windows of development has yet to be fully elucidated.

Multiple cell types make up the liver with each cell type having specialized roles in metabolism, immunity, and tissue maintenance. Hepatocytes are the main functional cells important for xenobiotic biotransformation (Gumucio and Miller, 1981; Andrews *et al*., 2022). Cholangiocytes are important for bile acid modification and transport (Ishibashi *et al*., 2009). Endothelial cells mediate exchange between blood and liver cell types (Zhang *et al*., 2023). Stellate cells store vitamin A, regulate extracellular matrix production, and transform into hepatic myofibroblasts upon activation and as a response to inflammatory signals (MacParland *et al*., 2018; Andrews *et al*., 2022; Zhang *et al*., 2023). Kupffer cells are liver resident macrophages involved in hepatic immune surveillance (Nemeth *et al*., 2009). Other immune cell populations that reside in and circulate around the liver are also present at steady state, including B cells, T cells, natural killer (NK) cells, conventional and plasmacytoid dendritic cells (cDC and pDC, respectively), monocytes and monocyte-derived macrophages (MDM), neutrophils, and mast cells (Nemeth *et al*., 2009; MacParland *et al*., 2018; Andrews *et al*., 2022). Understanding the changes of PCB exposure during critical developmental time windows at the single cell level is important in elucidating the cell type-specific responses and interactions that can provide mechanistic insight. Therefore, in the present study, we performed single cell RNA sequencing in the offsprings that were perinatally exposed to PCBs. We hypothesized that maternal PCB exposure leads to cell type-specific responses involved in metabolic dysregulation in livers of the offspring.

## MATERIALS AND METHODS

### Chemicals

The Fox River PCB mixture was prepared from technical PCB mixtures. Aroclor 1242, 1248, 1254, and 1260 were prepared in pesticide grade acetone (50 mg/ml acetone) (purchased from Fisher Scientific, Far Lawn, New Jersey) and mixed at a ratio of 35:35:15:15, respectively, by weight. The acetone was then evaporated, and the solute PCB mixture was used in the study upon successful congener-specific analysis. Full details of the PCB composition and mixture generation methods are described in(Kostyniak *et al*., 2005). The PCB mixture was dissolved in organic peanut oil (Spectrum Organic Products, LLC, Melville, NY) at a stock concentration of 20 mg/ml and stored in amber glass vials at − 20 ◦C.

### Animals and chemical exposure

All mice were housed according to the Association for Assessment and Accreditation of Laboratory Animal Care International guidelines (https://aaalac.org/resources/theguide.cfm). All experiments were approved by the Institutional Animal Care and Use Committee (IACUC) at the University of Washington. Three-month-old SPF-free female C57BL/6J mice were purchased from the Jackson Laboratory (Bar Harbor, Maine) and were acclimated to the animal facility at the University of Washington at 74 degrees Fahrenheit, 26% humidity, and 12-h light and dark cycle for 1 week prior to the experiment. All mice had ad libitum access to standard laboratory autoclaved rodent diet (LabDiet No. 5010 after weaning and LabDiet No. 5021 for breeding; LabDiet, St Louis, Missouri) and water (non-acidified autoclaved water). Animals were housed with autoclaved bedding (autoclaved Enrich-N’Pure; Andersons, Maumee, Ohio). From 2 weeks before mating, female C57BL/6 mouse dams were exposed to organic peanut oil mixed with organic peanut butter (vehicle, Trader Joe’s, Monrovia, CA) or the Fox River PCB mixture in vehicle (6 mg/kg body weight) once daily until pups were weaned at postnatal day (PND) 21. Female mouse livers of the pups at PND 28 were collected. Female mice were used in this study to compare results with previous publications on PCB-mediated developmental neurotoxicity and as follow up to our previous work (Cheng *et al*., 2018; Lim *et al*., 2020; Li *et al*., 2022; Wilson *et al*., 2024). The experimental design is summarized in Fig. 1.

**Figure 1.**
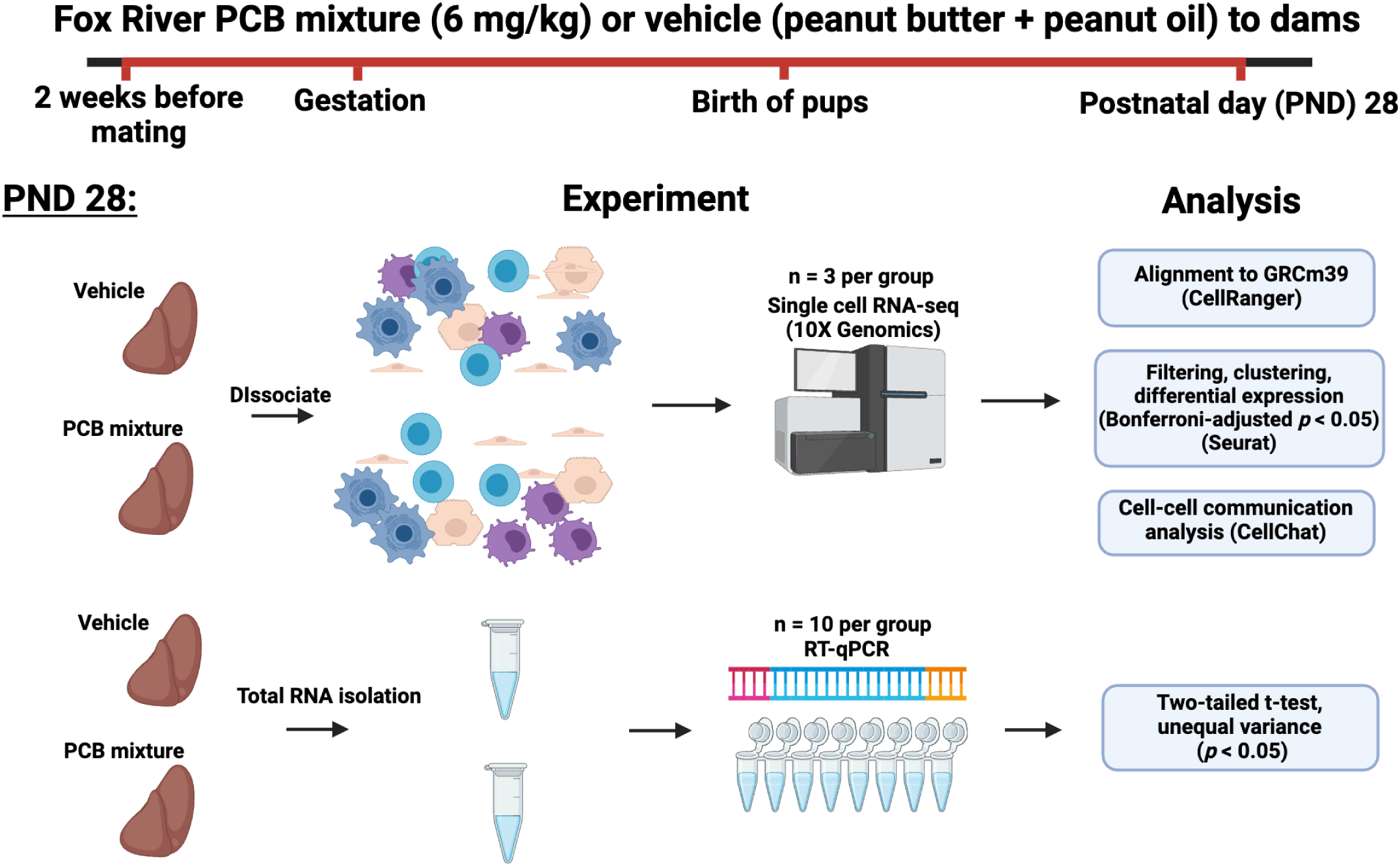
Experimental design of the study. Dams were exposed to peanut oil in peanut butter (vehicle) or the Fox River PCB mixture (6 mg/kg body weight) once daily until pups were weaned at PND 21. Using the livers of pups at PND 28, scRNA-seq (n = 3) was performed to profile the cell type-specific responses in liver or RT-qPCR (n = 10) to investigate whole liver mRNA expression changes.

### Liver dissociation

Fresh whole livers rinsed in Dulbecco’s phosphate-buffered saline (DPBS) (Cat No. 14190136, Thermo Fisher Scientific, Waltham, Massachusetts) and were minced to 1–3 mm pieces using surgical scissors (n=3 for each group). The minced livers were then placed in 15 ml conical tubes containing 10 ml of DPBS. Using serological pipettes, the DPBS was replaced by 10 ml of dissociation enzyme mixtures containing Liberase (Cat No. 540115001, Sigma Aldrich, St Louis, Missouri) and Dispase II (Cat No. D4693-1G, Sigma Aldrich, St Louis, Missouri) dissolved in Hanks’ Balanced Salt Solution (Cat No. 14025-092, Thermo Fisher Scientific, Waltham, Massachusetts). The tubes with the liver and enzyme mix were incubated at 37°C for 40 min using a Roto-Therm H2020 incubator (Benchmark Scientific Inc., Sayreville, New Jersey). After the incubation, the cells and undissociated fragments were strained using a 40-micron cell strainer. Strained cells were centrifuged at 300 × g for 5 min at 4°C and resuspended in a 5-ml red blood cell lysis buffer on ice for 8 min. Cells were centrifuged again at 300 × g for 5 min at 4°C. Dead and dying cells were filtered using the Dead Cell Removal Kit (Miltenyi Biotec, Cologne, Germany) following the manufacturer’s instructions. The viability was checked using a hemocytometer under a light microscope (Labophot-2, Nikon, Tokyo, Japan). Samples with >85% viability were then cryopreserved using 10% dimethylsulfoxide (Cat No. BP231-100, Thermo Fisher Scientific, Waltham, Massachusetts) and 90% fetal bovine serum (Cat No. F2442-50ML, Sigma Aldrich, St Louis, Missouri) until further analysis.

### Single cell RNA sequencing

Cryopreserved cells were thawed using a water bath at 37°C for 2 min, followed by serial dilution in DPBS until 32 ml was reached. Cells were centrifuged and resuspended in DPBS until a concentration of approximately 100 cells/μl was reached. The resuspended cells (n = 3), targeting 10 000 cells per sample, were then subject to scRNA-seq using a Chromium Next GEM single cell 3′ v3.1 kit and a Chromium X controller (10X Genomics, Pleasanton, California) following the manufacturer’s instructions. The created libraries were then sequenced using the NovaSeq platform.

### Data analysis of single cell RNA sequencing

Raw data were processed using the Cell Ranger v7.0 (10X Genomics, Pleasanton, California). Processed data were read into R version 4.2.2 (R Core Team, 2023) for further analyses. Filtering and normalization were performed using the default parameters using Seurat v5 (Hao et al., 2021). All cells with number of genes < 200 and total reads < 20,000 and mitochondrial RNA percent < 40 were filtered out to account for low-quality cells, doublets, and dying cells, respectively. Clustering was performed using the first 50 principal components. Cell clusters between vehicle and PCB exposed groups were integrated prior to cell type labeling using Harmony(Korsunsky *et al*., 2019). Cell type labeling was performed using the differentially expressed genes using the function FindAllMarkers with default parameters in Seurat (Bonferroni-adjusted p-value < 0.05). Cell-cell communication analysis was done using the CellChat (v. 1.5) package (Jin *et al*., 2021). The intercellular communication results were selected based on significantly enriched predicted signaling with an information flow (sum of log probability) cutoff of 10. Gene ontology enrichment was done using the topGO (v.2.48.0) package(Adrian Alexa, 2017) using all detected genes as the background (FDR-adjusted p-value < 0.05). Heatmaps were created using the ComplexHeatmaps (v. 2.13.1) package(Gu *et al*., 2016; Gu, 2022). All other plots were created using ggplot2 (v. 3.3.6).

### Total RNA extraction and reverse transcription-quantitative polymerase chain reaction (RT-qPCR)

Total RNA was isolated from the large intestine using RNA-Bee (Tel-Test Inc., Friendswood, Texas). RNA concentrations were quantified using a NanoDrop 1000 Spectrophotometer (Thermo Scientific, Waltham, Massachusetts) at 260 nm. The integrity of total RNA samples was evaluated by formaldehyde-agarose gel electrophoresis with visualization of 18S and 28S rRNA bands under UV light. Extracted RNA samples were then reverse transcribed to cDNA using a High-Capacity cDNA Reverse Transcription Kit (Life Technologies, California). The resulting cDNA products were amplified by qPCR, using a Sso Advanced Universal SYBR Green Supermix in a Bio-Rad CFX384 Real-Time PCR Detection System (Bio-Rad, Hercules, California). Data were normalized to the housekeeping gene Glyceraldehyde 3-phosphate dehydrogenase (Gapdh) using the ΔΔCq method and were expressed as % of Gapdh and p-value < 0.05 using t-test was set as the statistical significance threshold. Primer sequences are shown in Supplementary Table S1.

## RESULTS

### Cell type specific responses following perinatal exposure to PCBs

We utilized scRNA-seq to profile changes in cell type-specific signatures following perinatal exposure to PCBs. The clusters of liver cells representing the major cell types are shown in Fig. 2A. These cell types identified through scRNA-seq include 1) resident liver cell types (i.e., hepatocytes, cholangiocytes, endothelial cells, stellate cells, myofibroblasts, and Kupffer cells) as well as 2) circulating immune cell types that reach the liver (e.g., B cells, T cells, natural killer [NK] cells, monocyte-derived macrophages [MDM], conventional and plasmacytoid dendritic cells [cDC and pDC, respectively], and mast cells). Cell types were labeled using established marker genes that are uniquely expressed in each cell type that are widely used in other scRNA-seq analysis pipelines (Guilliams *et al*., 2022; Lim *et al*., 2024, 2025). Single cell clusters displayed minimal differences between vehicle and PCB exposure (Fig. S1). Each marker gene was enriched and was highly expressed in the corresponding liver cell type (Fig. 2B).

**Figure 2.**
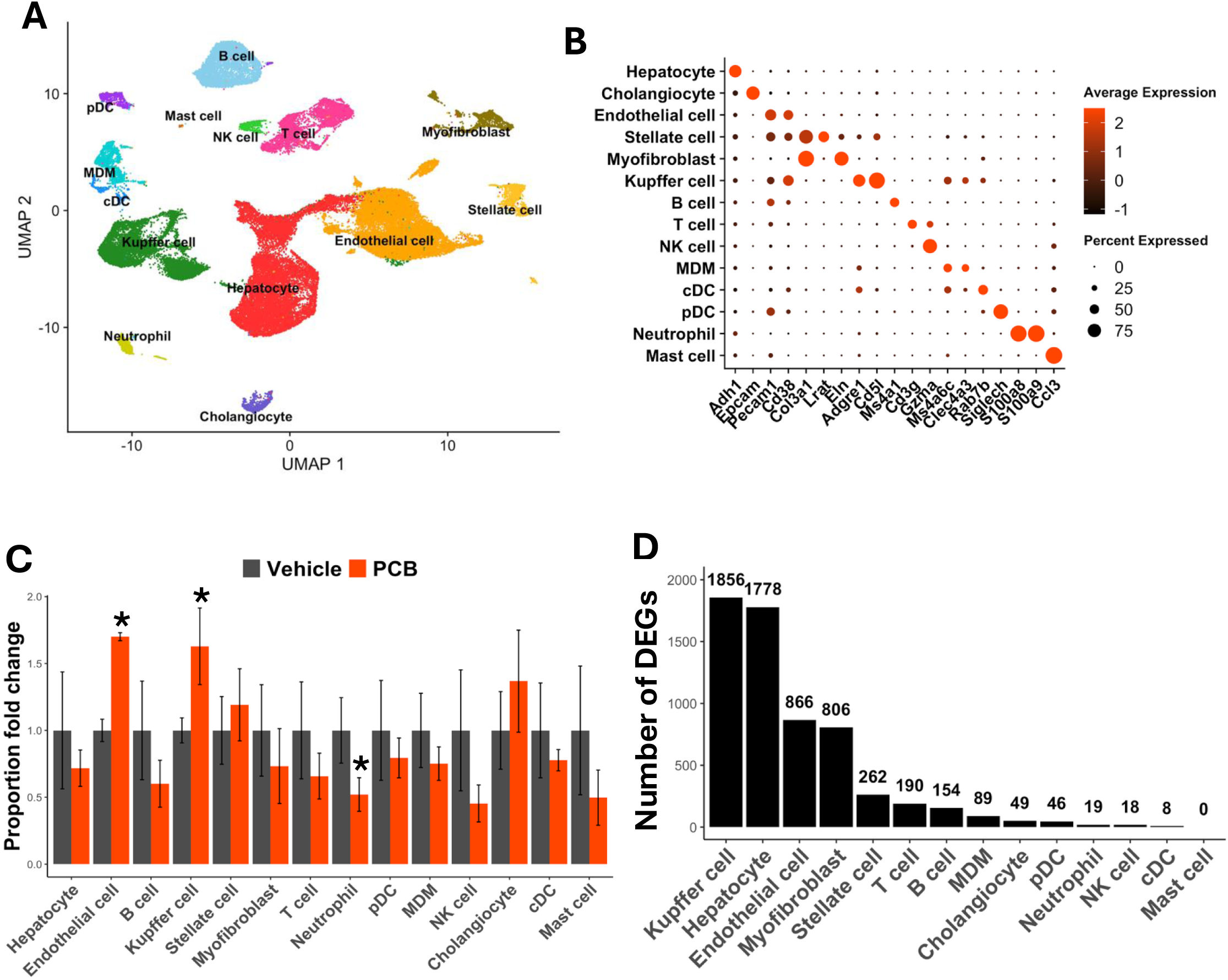
Clustering and visualization of marker genes in each cell type of the liver. **A.** Visualization of cell clusters with cell types identified using marker gene expression. **B.** Representation of key marker genes for cell type labeling. **C.** Changes in proportions of liver cell types following early life exposure to PCBs. Y-axis shows the fold change in each cell type of the PCB exposed group over that of the vehicle exposed group. Asterisks represent p-value (two-way t-test with unequal variance assumption). **D.** Total number of differentially expressed genes (Bonferroni-adjusted *p*-value < 0.05) for each liver cell type from perinatal exposure to PCBs.

Each hepatic cell type performs specialized functions, and certain chemicals may preferentially impact specific cell populations (Laskin, 1990; Siwicki *et al*., 2021; Lim *et al*., 2024). To determine whether early PCB exposure alters liver cellular composition, we analyzed cell type proportions using scRNA-seq. We observed a statistically significant increase in endothelial cell and Kupffer cell populations alongside a significant decrease in neutrophil populations, in PCB-exposed offsprings compared to vehicle-exposed controls (t-test with unequal variance *p* < 0.05, Fig. 2C).

To characterize Cell type-specific transcriptomic alterations by early life PCB exposure, we performed differential expression analysis for each liver cell type (Bonferroni-adjusted *p*-value < 0.05) (Fig. 2D and Table S2). Overall, PCBs mostly impacted the transcriptomic signatures in resident liver cell types over other immune cell types. Among resident cells, Kupffer cells exhibited the highest number of differentially expressed genes (DEGs), followed by hepatocytes, endothelial cells, and myofibroblasts.

To gain further insights into cell type-specific transcriptomic alterations by early life PCB exposure, we performed gene ontology enrichment analysis on the differentially expressed genes (DEGs) identified in each cell (Fig. 3). The four cell types with the highest number of DEGs exhibited distinct response patterns following early life PCB exposure. In Kupffer cells, genes associated with cell migration, cytoskeleton organization, and signal transduction were up-regulated, whereas genes related to endocrine protein response and immune response were down-regulated (Fig. 3A); in hepatocytes, genes involved in aerobic energy were up-regulated, whereas genes related to xenobiotic biotransformation and peptide hormone response were down-regulated (Fig. 3B); Endothelial cells displayed up-regulation of genes involved in cell migration and vascular endothelial growth factor signaling, alongside down-regulation of interferon signaling and innate immune response-related genes (Fig. 3C). In myofibroblasts, genes linked to cellular lipid efflux were up-regulated, whereas those associated with endoplasmic reticulum stress were down-regulated (Fig. 3D).

**Figure 3.**
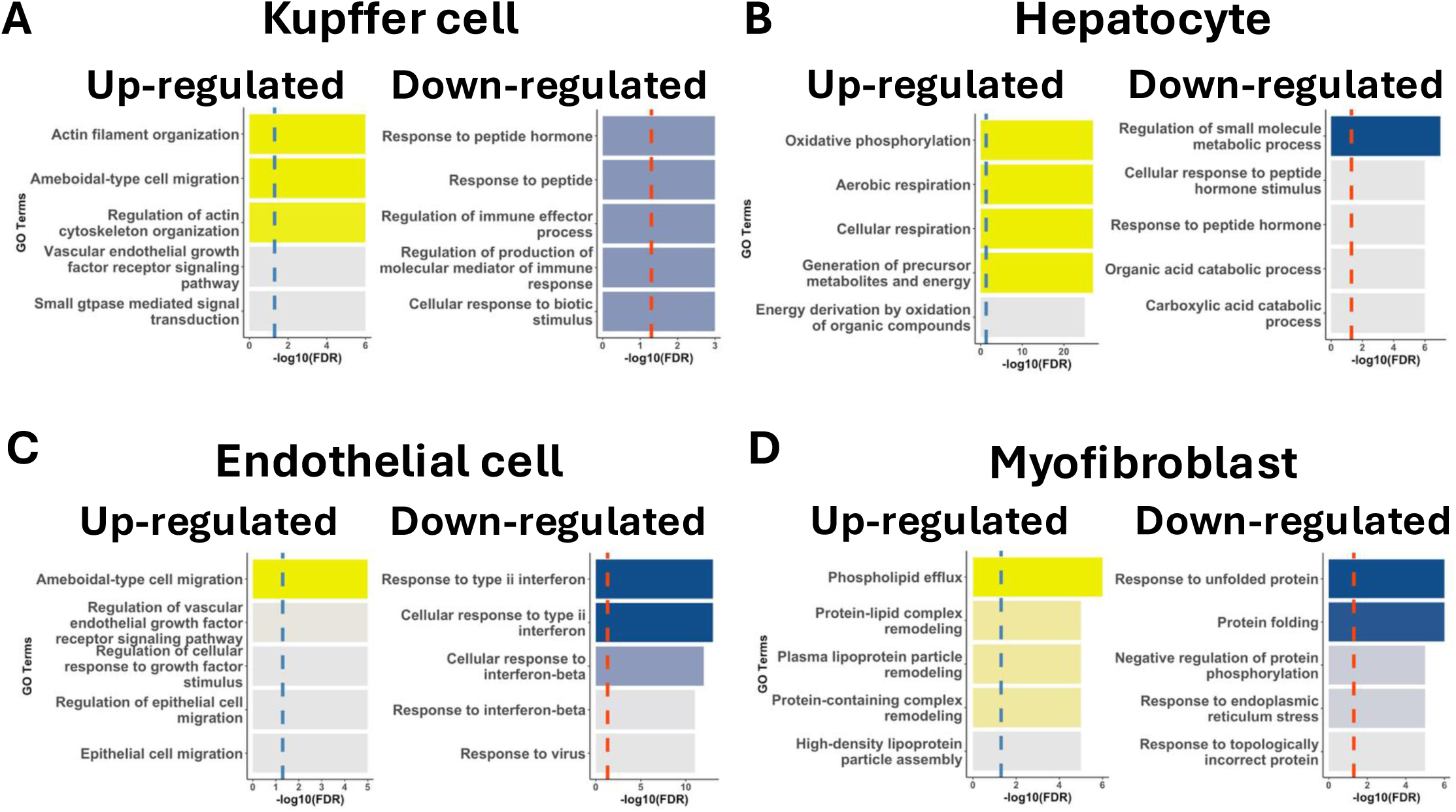
Top 5 up- and down-regulated gene ontology enrichment results in Kupffer cell (**A**), hepatocyte (**B**), endothelial cell (**C**), and myofibroblast (**D**) from perinatal exposure to PCBs. Dotted lines represent -log10 FDR-adjusted p-value at 0.05.

In other liver cell types, early life PCB exposure produced distinct transcriptomic changes across various biological functions. In stellate cells, genes involved in growth factor response and angiogenesis were up-regulated, whereas those related to xenobiotic and sterol metabolism were down-regulated (Fig. S2A). In cholangiocytes, genes associated with cellular lipid transport and clearance were up-regulated (Fig. S2B). In monocyte-derived macrophages (MDMs), genes involved in the regulation of hematopoiesis and cell adhesion, and metal ion response were up-regulated, whereas chemotaxis-related genes were down-regulated (Fig. S2C). In B cells, T cells, and NK cells, genes involved in fatty acid and xenobiotic metabolism were up-regulated (Fig. S3A and Fig. S3C). In both conventional and plasmacytoid dendritic cells (cDC and pDC), genes related to sterol and lipid transport and metabolism were up-regulated (Fig. S3D-S3E). Collectively, these findings demonstrate that the hepatic transcriptomic response to early life PCB exposure is highly cell type-specific.

### Upregulation of major drug processing genes following early life exposure to PCBs

One major function of the liver is xenobiotic biotransformation through drug processing genes that encode for enzymes in phase-I and phase-II metabolism as well as transporters (Almazroo *et al*., 2017). We and others previously showed that both acute and early life exposure to environmental toxicants dysregulate the hepatic transcriptome and functions (Kim *et al*., 2014, 2023; Lim *et al*., 2021, 2024). Furthermore, we recently showed that in addition to hepatocytes, the gene expression landscape varies within hepatic cell types, with non-parenchymal cells expressing unique portfolios of drug processing genes (Lim *et al*., 2024, 2025). Therefore, we further investigated the cell type-specific changes in drug processing gene expression following early life PCB exposure.

Coplanar (dioxin-like) PCBs are known to activate the aryl hydrocarbon receptor (AhR) with cytochrome P450 (*Cyp*) *1a2* as the prototypical target (Bemis *et al*., 2005). In contrast, noncoplanar (non-dioxin like) PCB congeners activate constitutive androstane receptor (CAR) and pregnane X receptor (PXR), with *Cyp2b10* and *Cyp3a11* as their prototypical targets, respectively(Al-Salman and Plant, 2012). Fox River PCB mixture contains both coplanar and noncoplanar PCB congeners (Kostyniak *et al*., 2005). In addition, some PCB congeners have been shown to activate the lipid-sensing nuclear receptor peroxisome proliferator-activated receptor alpha (PPARα) (Wahlang *et al*., 2014; Routti *et al*., 2019), with Cyp4a10 being a prototypical target gene (Rakhshandehroo *et al*., 2010). Consistent with this, early life exposure to the Fox River PCB mixture up-regulated the prototypical target genes of AhR, PXR, CAR, and PPARα in hepatocytes (Fig. 4A).

**Figure 4.**
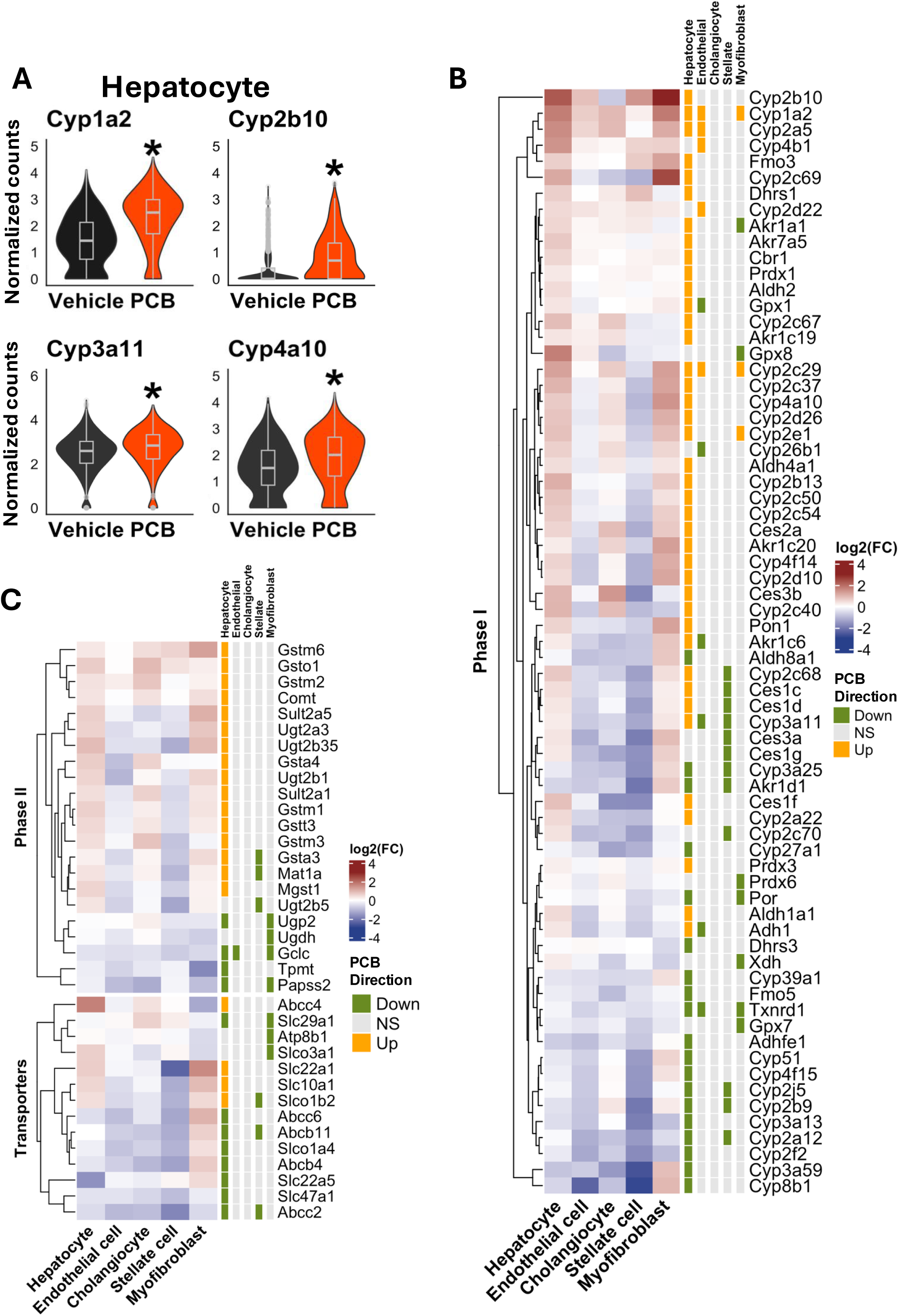
Dysregulated expression signatures of drug-processing genes in resident hepatic cell populations following perinatal exposure to PCBs. **A.** Violin plots that describe the distribution of expression of prototypical target genes of aryl hydrocarbon receptor (AHR), pregnane X receptor (PXR), constitutive androstane receptor (CAR), and peroxisome proliferator-activated receptor alpha (PPARα) from perinatal exposure to PCBs. Asterisks represent Bonferroni-adjusted p-value < 0.05. **B.** Phase-I drug-processing enzymes that were differentially expressed (rows) in at least one of the major hepatic resident cell types (columns) with xenobiotic biotransformation capabilities are shown in a heatmap (left side). The colors of the heatmap represent the log2 fold change of liver genes of the PCB exposed pups as compared to the vehicle control. “Direction” indicates whether a gene is up-(orange) or down-(green) regulated from perinatal exposure to PCBs (Bonferroni-adjusted p-value < 0.05). **C.** Phase-II enzymes and transporters involved in xenobiotic metabolism processes that were differentially expressed (rows) in at least one of the major hepatic resident cell types (columns) with xenobiotic biotransformation capabilities are shown in a heatmap (left side). The colors of the heatmap represent the log2 fold change of liver genes of the PCB exposed pups as compared to the vehicle control. “Direction” indicates whether a gene is up-(orange) or down-(green) regulated from perinatal exposure to PCBs (Bonferroni-adjusted p-value < 0.05).

Early life PCB exposure broadly dysregulated drug processing genes across multiple liver cell types. In hepatocytes, most differentially expressed phase-I enzymes were up-regulated by early life PCB exposure (Fig. 4B), including those in the *Cyp2*, aldo-keto reductases (*Akr*), aldehyde dehydrogenases (*Aldh*), and carboxylesterases (*Ces*) families. Conversely, many phase-II enzymes were down-regulated in hepatocytes (Fig. 4C), despite some up-regulation in glutathione-S transferase (*Gst*), sulfotransferase (*Sult*), and UDP glucuronosyltransferase (*Ugt*) families. Notably, key synthetases for phase-II enzyme cofactors, including UDP-glucose pyrophosphorylase 2 (*Ugp2*, for synthesizing UDP-glucuronic acid), glutamate-cysteine ligase catalytic subunit (*Gclc*, for synthesizing glutathione), and 3’-Phosphoadenosine 5’-Phosphosulfate Synthase 2 (*Papss2*, for synthesizing PAPS), where down-regulated, potentially limiting conjugation capacity.

Transporters expression was also dysregulated by early life PCB exposure in hepatocytes. Most up-regulated transporters were basolateral uptake transporters, including solute carrier family (*Slc*) *10a1* (sodium-taurocholate cotransporting polypeptide, Ntcp), *Slc22a1* (organic cation transporter 1, Oct1), and solute carrier organic anion transporter (*Slco*) *1b2* (organic anion transporting polypeptide 1b2, Oatp1b2). In contrast, except for ATP-binding cassette (*Abc*) *c4* (multidrug resistance protein 4, Mrp4), which was upregulated by PCBs, most down-regulated transporters were efflux transporters, including *Abcc2* (Mrp2), *Abcc6* (Mrp6), *Abcb11* (Bsep), *Abcb4* (Mdr2), and *Slc47a1* (Mate1). The bi-directional equilibrative nucleoside transporter 1 (Ent1/Slc29a1), as well as the uptake transporters of *Slc22a5* (novel organic cation transporter, Octn2) and *Slco1a4* (Oatp1a4), were down-regulated by PCBs.

In addition to hepatocytes, early life exposure to PCBs led to sparse but notable dysregulation of drug-processing genes in multiple resident liver cell types, including endothelial cells, cholangiocytes, stellate cells, and myofibroblasts. Most phase-I metabolism genes were up-regulated in hepatocytes but down-regulated in non-parenchymal cells (Fig. 4B). Notably, some genes that were up-regulated in hepatocytes, such as members of the *Ces* and *Cyp* families, *Gpx1*, and *Adh1*, were concurrently down-regulated in non-parenchymal cells. Similarly, *Gsta3* and *Mat1a* were up-regulated in hepatocytes but down-regulated in stellate cells. These opposing patterns suggest distinct cell type-specific transcriptional regulation by PCBs.

A subset of drug processing genes, including *Cyp4b1*, *Cyp26b1*, *Akr1a1*, *Ugdh*, *Papss2*, *Slco3a1*, and *Slc29a1*, exhibited higher expression in non-parenchymal cells compared to hepatocytes under basal conditions (Fig. S4), consistent with our previous characterization on basal expression patterns in liver cell types (Lim *et al*., 2025). However, the majority of dysregulated drug processing genes, in both control and PCB-exposed groups, were most highly expressed in hepatocytes (Fig. S4), suggesting that non-parenchymal cells are unlikely to fully compensate for hepatocyte-specific dysregulation.

### Downregulation of insulin signaling and glucose metabolism-related genes from early life exposure to PCBs

Interestingly, of all liver cell types detected, most resident liver cell types, except cholangiocytes, exhibited down-regulated responses related to insulin signaling and glucose metabolism. Notably, a subset of genes involved in glucose response was up-regulated in hepatocytes (Fig. 5A and Table S3). Genes involved in insulin stimulus response were down-regulated in both hepatocytes and Kupffer cells, whereas genes involved in insulin receptor signaling were down-regulated in hepatocytes and myofibroblasts. In endothelial cells, genes involved in glucose storage and transport were down-regulated, along with hepatocytes and endothelial cells both exhibiting decreased expression of genes involved in glucose homeostasis. In addition, genes related to glucose metabolism were down-regulated in both hepatocytes and stellate cells. Other down-regulated gene ontology terms in hepatocytes include insulin receptor signaling and intracellular glucose homeostasis.

**Figure 5.**
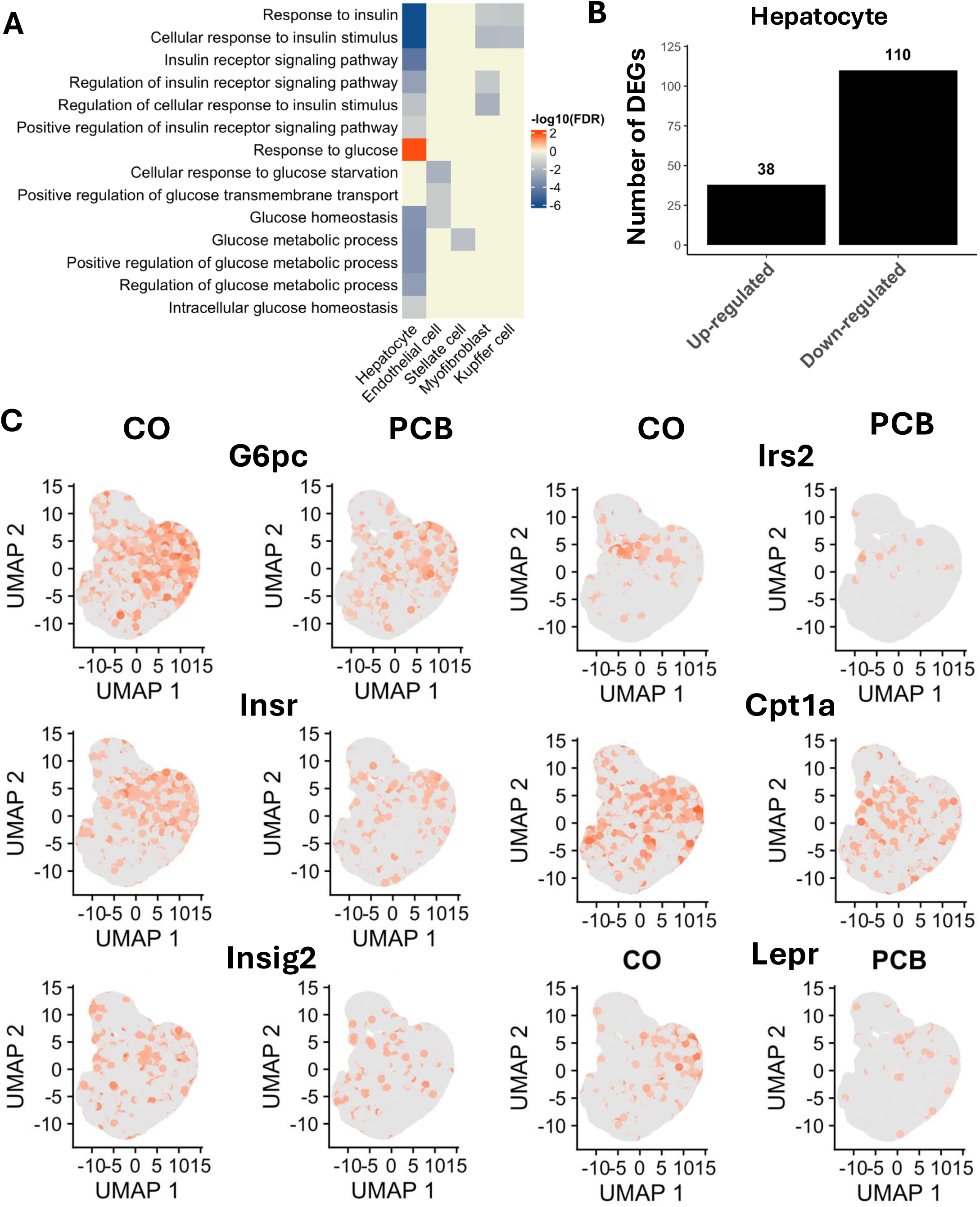
A. Representation of up-regulated cell type-specific gene ontology terms related to insulin signaling and glucose homeostasis from perinatal PCB exposure. Red and blue show up- and down-regulated gene ontology terms, respectively. Color gradient indicates -log10 transformed FDR-adjusted p-value. **B**. Number of up- and down-regulated genes related to glucose and insulin signaling in hepatocytes. **C**. Down-regulated expression of glucose and insulin signaling markers in hepatocytes. Grey and red indicate low and high expression, respectively.

Overall, the majority of down-regulated genes related to insulin signaling and glucose metabolism were found in hepatocytes (Fig. 5A), suggesting that hepatocytes are the primary targets of metabolic disruption following early life PCB exposure. To determine the overall directionality of these pathway signatures, we compared the number of up-regulated vs. down-regulated genes involved in insulin and glucose metabolism in hepatocytes. Our findings showed a greater number of down-regulated genes than up-regulated genes (Fig. 5B). Representative examples of key down-regulated genes are shown in Fig. 5C. These include glucose 6-phosphatase (*G6pc*), a key enzyme in glucose metabolism during gluconeogenesis and glycogenolysis (Marcolongo *et al*., 2013), insulin receptor substrate 2 (*Irs2*), which mediates hepatic insulin signaling and regulates glucose and lipid metabolism(Geng *et al*., 2014), and Insulin receptor (*Insr*) (Lee and Pilch, 1994). Other notable genes include carnitine palmitoyltransferase 1A (*Cpt1a*), a key regulatory of fatty acid import for hepatic β-oxidation(Schlaepfer and Joshi, 2020), insulin-induced gene 2 (*Insig2*), which inhibits sterol regulatory element binding protein (SREBP) activation, thereby suppressing lipid biosynthesis (Yabe *et al*., 2003), and leptin receptor (*Lexpr*), which mediates leptin signaling and influences insulin sensitivity (Polyzos *et al*., 2015). In summary, these findings indicate that early life PCB exposure induces metabolic dysfunction in hepatocytes, as evidenced by the predominant down-regulation of key genes involved in insulin signaling and glucose homeostasis.

### Upregulation of endoplasmic reticulum (ER) stress-related responses from early life exposure to PCBs

In hepatocytes, early life exposure to PCBs up-regulated genes involved in ER stress. Specifically, genes related to both the positive and negative regulation of ER stress, as well as ER stress-induced apoptosis were up-regulated (Fig. 6A). In contrast, genes related to ER stress, unfolded protein response, topologically incorrect protein response, inositol-requiring transmembrane kinase/endonuclease (Ire1)-mediated unfolded protein response were down-regulated in endothelial cells and myofibroblasts. Meanwhile, in plasmacytoid dendritic cells (pDCs) and T cells, genes involved in protein folding and protein refolding were up-regulated from early life exposure to PCBs (Fig. 6A).

**Figure 6.**
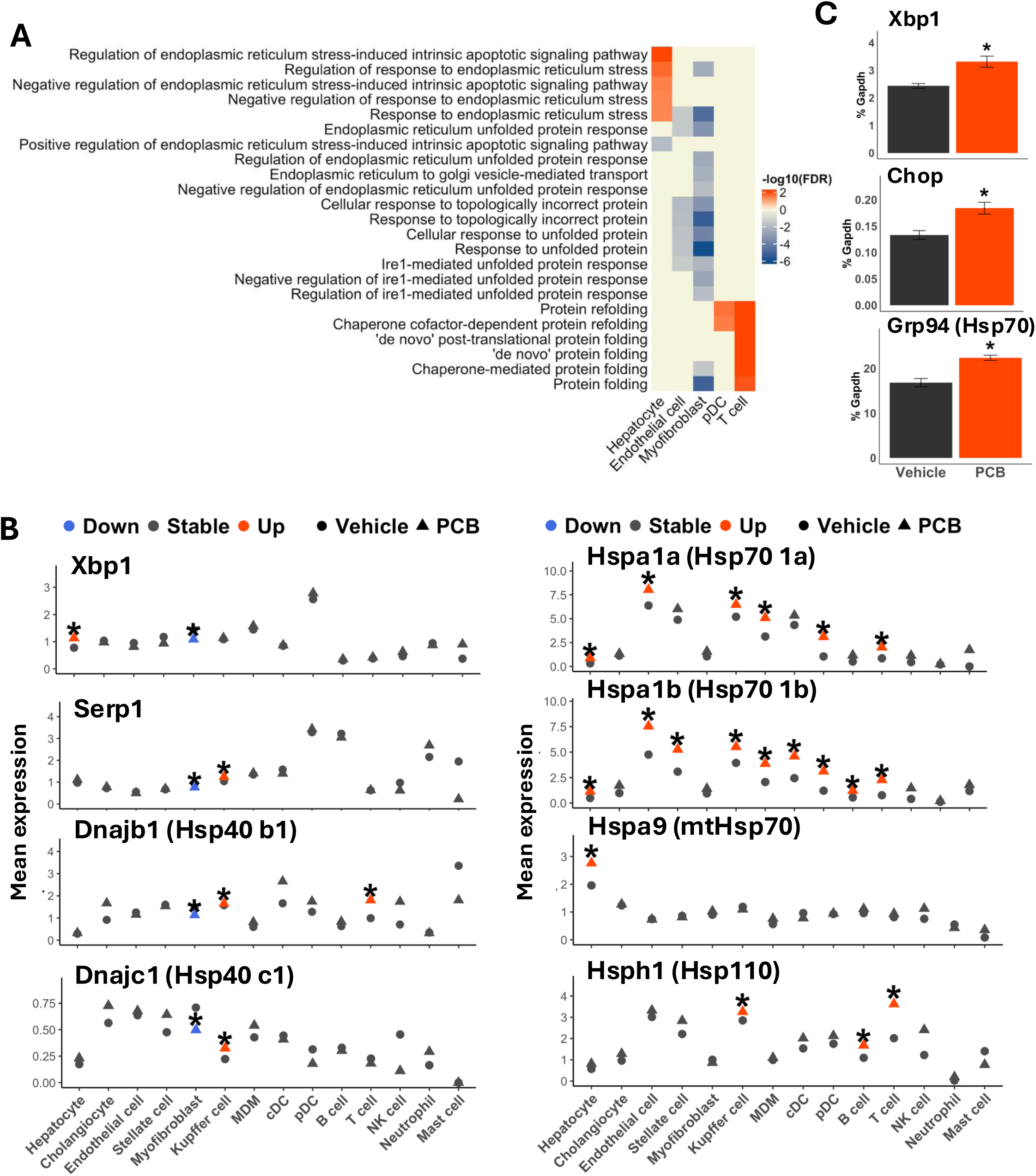
**A**. Representation of up-regulated cell type-specific gene ontology terms related to insulin signaling and glucose homeostasis from perinatal PCB exposure. Red and blue show up- and down-regulated gene ontology terms, respectively. Color gradient indicates -log10 transformed FDR-adjusted p-value. **B**. Average expression of endoplasmic reticulum stress markers in hepatic cell types following perinatal PCB exposure. Red and blue colors represent up- and down-regulation, respectively. Vehicle and PCB-exposed groups are shown as circles and triangles, respectively. Asterisks indicate differential expression (Bonferroni-adjusted p-value < 0.05). **C**. Up-regulated mRNA expression of endoplasmic reticulum stress markers by RT-qPCR in livers perinatally exposed to PCBs. Asterisks show p-value < 0.05 (two-way t test with assumption of unequal variance).

Multiple components of ER stress response were dysregulated by early life PCB exposure, including unfolded protein response (UPR), protein folding chaperones, and regulators. X-box binding protein 1 (Xbp1), a key transcription factor that is activated during ER stress(Park *et al*., 2021), was up-regulated in hepatocytes. Stress-associated endoplasmic reticulum protein 1 (Serp1), which supports protein folding and protects cells from ER stress-induced apoptosis (Hu *et al*., 1996), was up-regulated in Kupffer cells. DnaJ heat shock protein members B1 and C1 (Dnajb1 and Dnajc1, respectively) are members of the heat shock protein 40 (Hsp40), which facilitates ER-associated degradation (ERAD) and recruits the immunoglobulin binding protein (BiP) during ER stress (Chen *et al*., 2017; Pobre *et al*., 2019). Hsp family A members 1a and 1b (Hspa1a and Hspa1b, respectively) encode Hsp70, which binds misfolded proteins during ER stress (Chen *et al*., 2022). Hsp family A member 9 (Hspa9) is a mitochondrial Hsp (mtHsp70) that coordinates the mitochondrial response to ER stress (Gorrell *et al*., 2022). Hsp family H member 1 (Hsph1, also known as Hsp110) enhances the refolding capacity of Hsp70 and stabilizes denatured proteins (Easton *et al*., 2000). Overall, early life PCB exposure up-regulated ER stress markers in multiple cell types, with the exception of myofibroblasts (Fig. 6B). Xbp1 was up-regulated in hepatocytes, whereas Serp1 was up-regulated in Kupffer cells.

Regarding the mRNA regulation of heat shock proteins, which are essential components of ER stress response, Hsp40 members b1 (Hsp40 b1) was down-regulated in myofibroblasts and up-regulated in Kupffer cells and T cells; and Hsp40 member c1 (Hsp40 c1) showed a similar pattern, which is down-regulation in myofibroblasts and up-regulated in Kupffer cells. In addition, Hsp70 member 1a (Hsp70 1a) was up-regulated in hepatocytes, endothelial cell, Kupffer cells, MDMs, pDCs, and T cells. Hsp70 member 1b (Hsp70 1b) was similarly up-regulated in these cell types, with the addition of B cells. Mitochondrial Hsp70 was up-regulated in hepatocytes, whereas Hsp110 was up-regulated in Kupffer cells, B cells, and T cells.

Consistent with the single-cell sequencing results, RT-qPCR using whole liver homogenates confirmed that Xbp1, C/EBP homologous protein (Chop), and Hsp70 transcripts were up-regulated from early life PCB exposure. Collectively, these results suggest that early life PCB exposure induces a robust ER stress response in the liver.

### Dysregulation of intercellular communication signaling pathways following early life PCB exposure

Multiple cell types in the liver were targeted from early life PCB exposure to varying degrees. To further investigate how these cell-type specific responses may contribute to altered liver pathophysiology, we performed ligand-receptor centered intercellular communication analysis. This approach identified 9 signaling pathways with significant interaction elements were enriched and predicted to be dysregulated from early life exposure to PCBs (Fig. 7A, Fig. S5, and Fig. S6), suggesting a shift in liver intercellular communication and physiology. The following signaling pathways were predicted to be down-regulated following early life PCB exposure: 1) Fibronectin (FN1), which is a critical component mediating cell adhesion, migration, and extracellular matrix remodeling (Zollinger and Smith, 2017); 2) laminin signaling, which supports the liver architecture and angiogenesis (Simon-Assmann *et al*., 2011; Goddi *et al*., 2021); 3) protease-activated receptors (PARs) signaling, which is involved in inflammatory crosstalk and vascular remodeling (Peach *et al*., 2023); 4) collagen signaling, which is essential for matrix support, cell migration, and fibrosis (Arteel and Naba, 2020; Zhang *et al*., 2022); 5) secreted phosphoprotein 1 (SPP1, Osteopontin) signaling, which interacts with integrins and functions as a chemotactic for macrophages (Han *et al*., 2023); 6) chemokine signaling pathways (CCL) signaling, which modulates the recruitment of immune cells, such as monocytes, T cells, and NK cells (Marra and Tacke, 2014); as well as 7) galectin signaling, which regulates cell death and immune modulation (An *et al*., 2021). 8) Amyloid precursor protein (APP) signaling in the liver, which mediates inflammatory response, oxidative stress, and cytokine release (Guo *et al*., 2021; Garcia *et al*., 2022), is predicted to be up-regulated by from early life PCB exposure. In addition, 9) CD45 (protein tyrosine phosphatase receptor, Ptprc) signaling, which is a marker of infiltrating lymphocytes, Kupffer cells, and monocytes and macrophages (Hermiston *et al*., 2003; De Simone *et al*., 2021), is predicted to be up-regulated by early life PCB exposure. In summary, early life exposure to PCBs dysregulated predicted signaling pathways that involve impaired immune activation and tissue remodeling.

**Figure 7.**
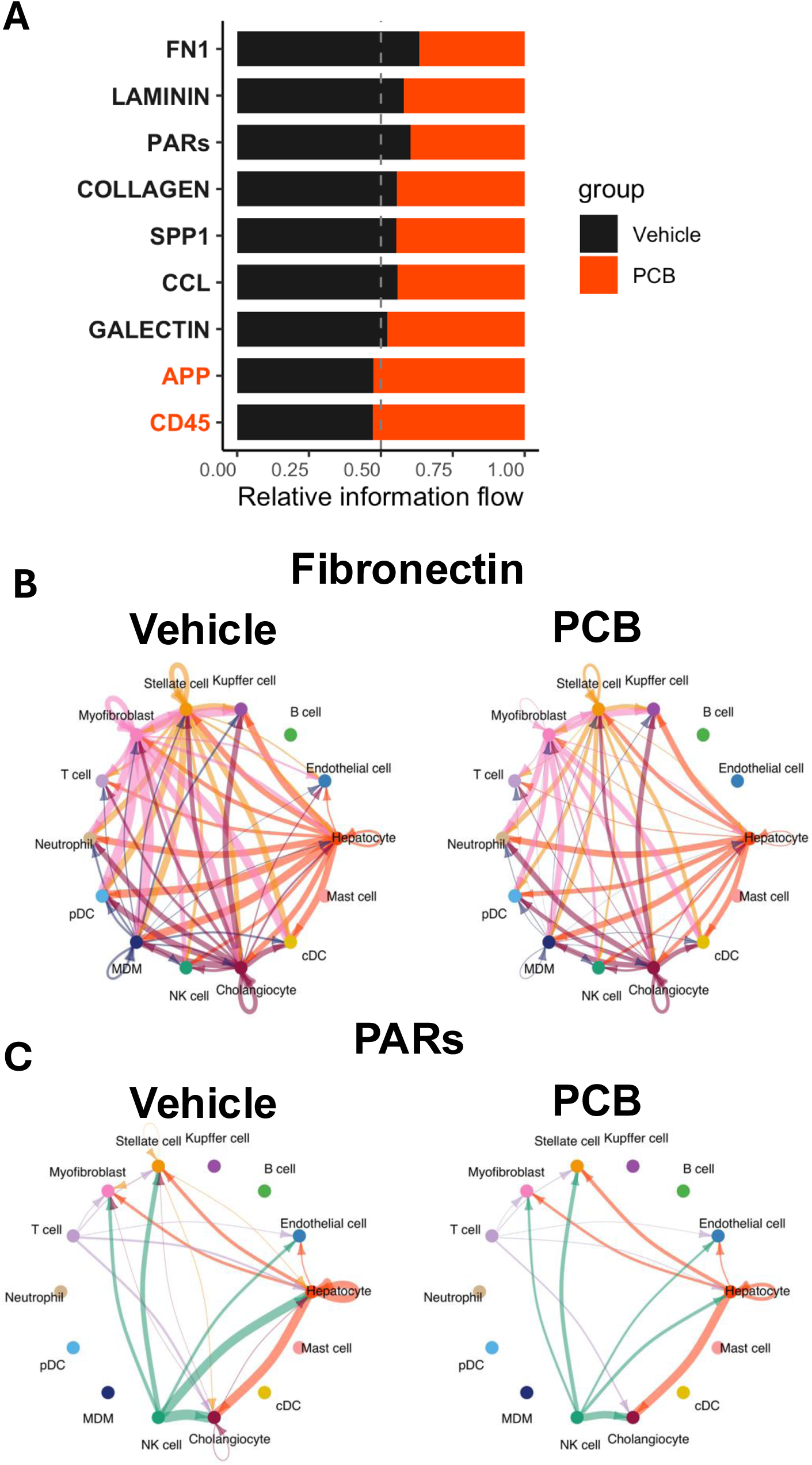
A. Significantly altered predicted cell-cell signaling pathways from perinatal PCB exposure (permutation test p-value < 0.05). Black and red highlights indicate significant predicted signaling pathways enriched in the vehicle and PCB exposed group, respectively. Predicted decrease in fibronectin (**B**) and increase in protease-activated receipts (PARs) (**C**) signaling in liver cell types from perinatal exposure to PCBs.

**Figure 8.**
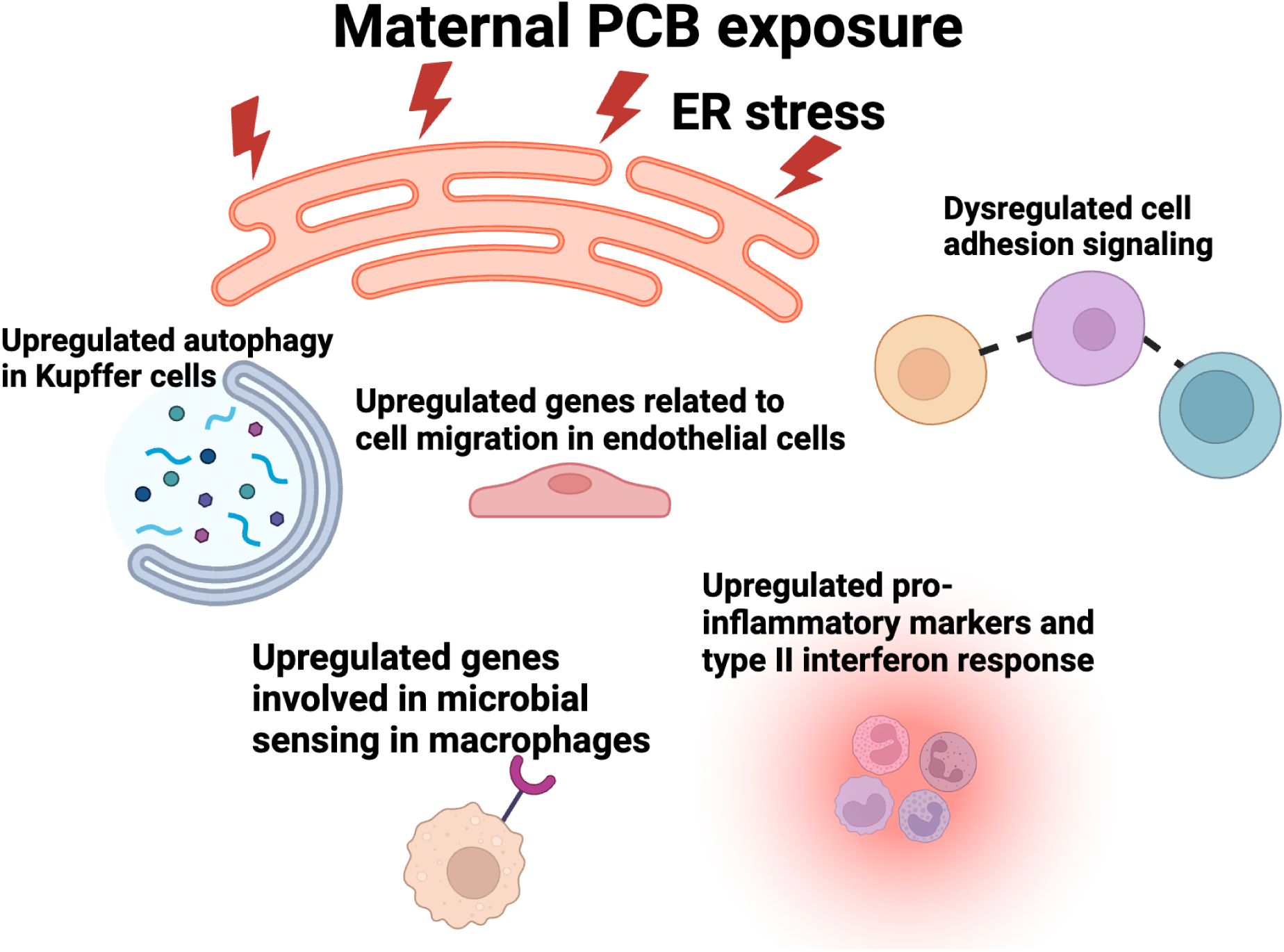
Summary of key findings.

## DISCUSSION

In the present study, we utilized single cell transcriptomics to investigate liver cell type-specific responses to early life exposure to Fox River PCB mixture in developing mouse liver. Our findings revealed that early life PCB exposure leads to transcriptional alterations in key pathways related to metabolic function, immune regulation, and intercellular communication. Among various cell types, Kupffer cells, hepatocytes, endothelial cells, and myofibroblasts exhibited the most prominent transcriptional changes, indicating they are the primary targets of PCB-induced hepatotoxicity.

Notably, early life PCB exposure dysregulated the expression of many genes involved in xenobiotic biotransformation, insulin signaling and regulation, and ER stress response in a cell type-specific manner. Our ligand-receptor interaction analysis further identified widespread alterations in intercellular communications. Our findings also suggest that dysregulated tissue homeostasis and immune signaling may be key mechanisms driving PCB-mediated hepatotoxicity.

Of the main cell types targeted by the early life PCB exposure, hepatocytes displayed broad dysregulation in genes involved in xenobiotic metabolism and transport. Early life PCB exposure up-regulated *Cyp1a2*, *Cyp2b10*, *Cyp3a11*, and *Cyp4a10*, which are the prototypical target genes for AhR (Vogel *et al*., 2020), CAR (Ding *et al*., 2006), PXR (Cui *et al*., 2010), and PPARα, respectively (Patsouris *et al*., 2006), in hepatocytes (Fig. 4), suggesting the activation of these transcription factors. In addition, many phase-I metabolism enzymes, including those in the *Ces*, *Akr*, *Gst*, and *Sult* families, were also up-regulated in hepatocytes by early life PCB exposure. Hepatocytes exhibited increased expression of many basolateral uptake transporters, including Oct1, Ntcp, and Oatp1b2, and decreased expression of many efflux transporters (Mrp2, Bsep, Mdr2, and Mate1). The up-regulation of key xenobiotic and lipid-sensing transcription factors and drug processing genes, as well as the coordinate regulation uptake and efflux transporters may serve as a compensatory mechanism to fight against PCB insult and contribute to altered kinetics of other xenobiotics from the environment.

One prominent observation was the PCB-mediated down-regulation of insulin signaling and glucose metabolism-related pathways in hepatocytes (Fig. 3 and Fig. 5). These findings align with previous studies showing PCB mixtures such as Aroclor 1254 impair key components of insulin signaling in mice (Zhang *et al*., 2015). Co-exposure to interesterified palm oil and PCB 126 exacerbated inflammation, morphological liver changes, and dysregulated carbohydrate metabolism in mice (Teixeira *et al*., 2024). PCB 126 dysregulated lipid metabolism and decreased polyunsaturated fatty acids in HepaRG cells (Mesnage *et al*., 2018). PCB exposure decreased hepatic glucose production and down-regulated genes involved in gluconeogenesis and glucose transport in rats and mice (Gray *et al*., 2013; Gadupudi *et al*., 2016). Furthermore, previous work reported that specific components of insulin signaling, and glucose metabolism were down-regulated by exposure to PCB exposure. For example, PCB 126 decreased hepatic *G6pc* gene expression in rats (Gadupudi *et al*., 2018), PCB 169 decreased hepatic expression of Cpt1a in mice (Wei *et al*., 2024), and circulating *LEPR* expression in children negatively correlated with PCB exposure levels in blood (Mondal *et al*., 2022). In line with previous findings, our results further support the disruption of insulin signaling and glucose metabolism by PCB exposure.

We and others have indicated that ER stress is likely a key mechanism underlying PCB-induced hepatotoxicity. We previously showed that exposure to the Fox River PCB mixture displayed ER stress-related transcriptomic signatures that are modulated by the microbiome in mice (Lim *et al*., 2020). Similarly, a single exposure to PCB 126 in rats up-regulated hepatic GRP78 at both the mRNA and protein levels, an effect associated with metabolic disruption (Chapados and Boucher, 2017). In HepG2 cells, PCB quinone, which is a PCB metabolite, up-regulated ER stress markers, including GRP78, GRP94, and CHOP, and induced ER structural abnormalities (Xu *et al*., 2015). Additionally, activation of the IRE1a-XBP1 pathway was observed following exposure to the PCB mixture (Arochlor 1254), which was associated with hepatic lipid accumulation in adolescent mice (Ruan *et al*., 2020). While key ER stress-related genes were up-regulated in the whole liver from maternal PCB exposure (Fig. 6B and Fig. 6C), hepatocytes displayed signatures related to unresolved or escalating ER stress, whereas myofibroblasts and endothelial cells have either resolved or suppressed UPR pathways as a potential compensatory mechanism to avoid apoptosis or to maintain vascular homeostasis. The different cell type-specific regulation of ER stress components from maternal PCB exposure suggests a temporal response to ER stress or a divergence in stress response mechanisms dependent on each cell type. Furthermore, ER stress is linked to down-regulated insulin signaling, altered energy and fatty acid metabolism (Lei *et al*., 2021; Lemmer *et al*., 2021; Park *et al*., 2022; Ma *et al*., 2024), which may overall exacerbate hepatotoxicity and metabolic dysfunction from PCB exposure.

We showed that non-parenchymal cells, particularly Kupffer cells and endothelial cells, are also primarily impacted by early life PCB exposure. PCB exposure increased proportions of Kupffer cells and endothelial cells, which displayed prominent transcriptomic responses as compared to other non-parenchymal cells (Fig. 2). Kupffer cells and endothelial cells showed up-regulated signatures of cell migration and down-regulated signatures involved in hepatic immune response (Fig. 3), suggesting tissue damage and matrix remodeling. These results were further supported by altered intercellular communication results involved in vascular remodeling, cell migration, and immune modulation targeting non-parenchymal cells (Fig. 7 and Fig. S6).

Our results align with previous work regarding PCB exposure and endothelial cell and Kupffer cell toxicity. Coplanar PCBs increase oxidative stress in endothelial cells and activate proinflammatory pathways that lead to cell dysfunction(Hennig *et al*., 2002). PCB exposure in mice fed a methionine-choline deficient diet (MCD) resulted in extrahepatic toxicity, including up-regulated circulating inflammatory biomarkers, suggesting endothelial cell dysfunction (Wahlang, Perkins, *et al*., 2017). PCB 126 induced endothelial cell inflammation that leads to increased expression of adhesion molecules (Han *et al*., 2012) as well as circulating levels of vascular inflammation markers that suggest endothelial cell dysfunction in mice (Wahlang, Barney, *et al*., 2017). PCB 126 polarized macrophages in vitro towards a proinflammatory phenotype and up-regulated chemokines that facilitate cell recruitment and infiltration (Wang *et al*., 2019). In rat Kupffer cells, chronic exposure to PCBs led to iron accumulation(Whysner and Wang, 2001), which is linked to increased oxidative stress and impaired function (Cui *et al*., 2022). In summary, our results, along with previous studies, suggest that maternal exposure to PCBs results in non-parenchymal cell injury, including endothelial cells and Kupffer cells, and suggest that non-parenchymal cells may play critical roles in modulating the hepatic response to PCB exposure.

The present study has several key limitations that warrant follow-up investigations on maternal PCB exposure and its effect on hepatic cell types in the developing offspring. The results of the present study are based on single cell transcriptomic data. Functional assays, such as western blot, metabolomics, glucose tolerance tests, and mitochondrial respiration assays, can help validate pathway-level conclusions on ER stress, altered insulin signaling and energy metabolism. Ligand-receptor interactions across cell types can be measured via additional experiments including spatial transcriptomics or secretome profiling. The present study investigated the cell type-specific responses shortly after perinatal PCB exposure in female mice. PCBs have been shown to disrupt endocrine signaling and display sex-specific outcomes (Wahlang *et al*., 2019; Liberman *et al*., 2020; Brennan *et al*., 2021). Follow up studies can use mice in both sexes to distinguish sex-specific vulnerability in hepatic cell type-specific responses following maternal PCB exposure. Furthermore, similar to previous studies (Lim *et al*., 2021, 2024; Kim *et al*., 2023), future work can assess whether altered hepatic cell type-specific responses persist into adulthood or lead to phenotypic changes later in life. Despite these limitations, our study has demonstrated that perinatal PCB exposure leads to cell type-specific transcriptomic alterations in the liver. Our findings highlight the multifaceted impact of early life environmental toxicant exposure on liver physiology and contribute to the understanding of developmental origins of metabolic and inflammatory liver disease.

