## Supplementary material for "Single-cell transcriptomics showed that maternal PCB exposure dysregulated ER stress-mediated cell type-specific responses in the liver of female offspring": PCB_single_cell_figures_draft_JL_V16_Supplemental.pdf

**Fig. S1**

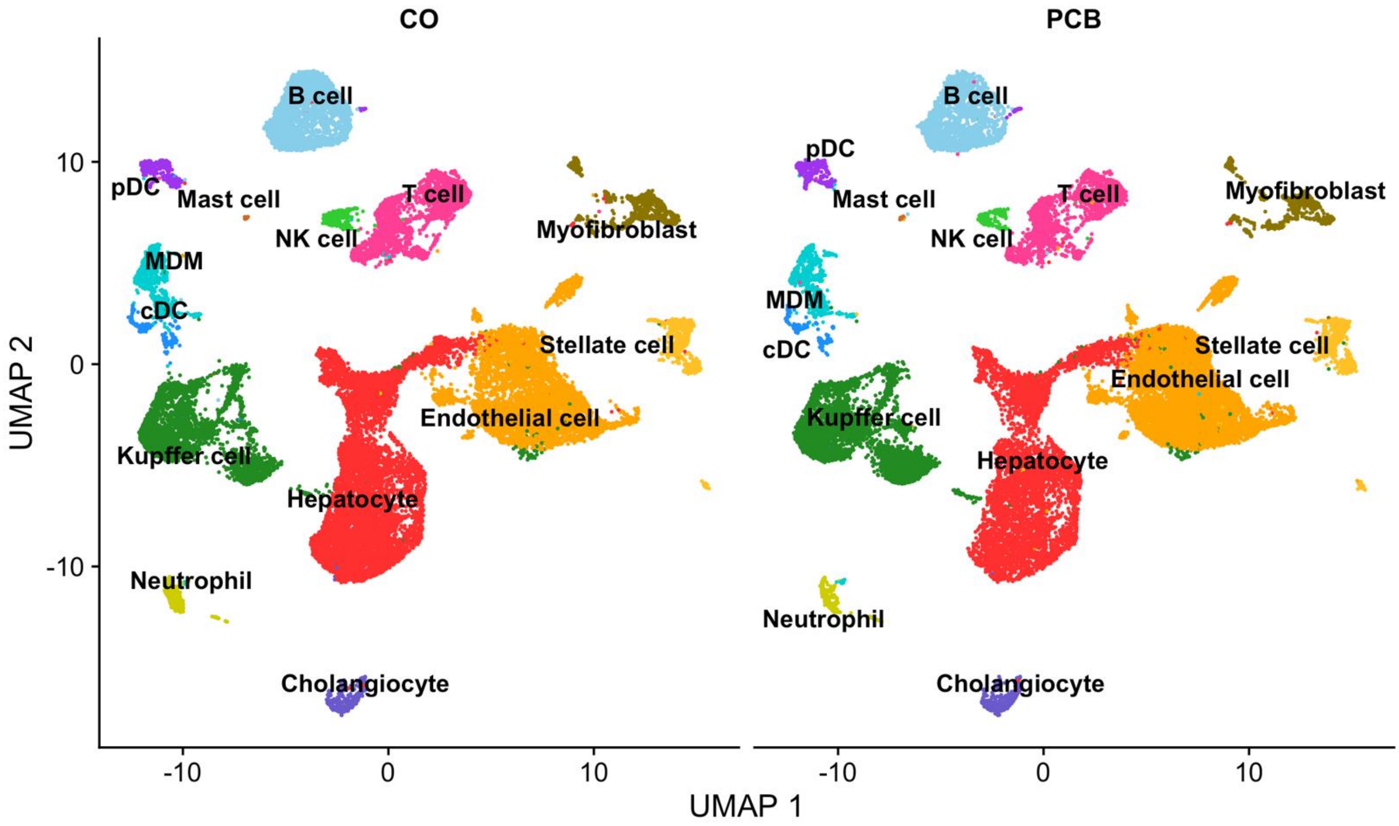


**Figure S1.** Labeled cell type clustering plot separated by early life exposure to vehicle (corn oil, CO) or PCB.

Fig. S2

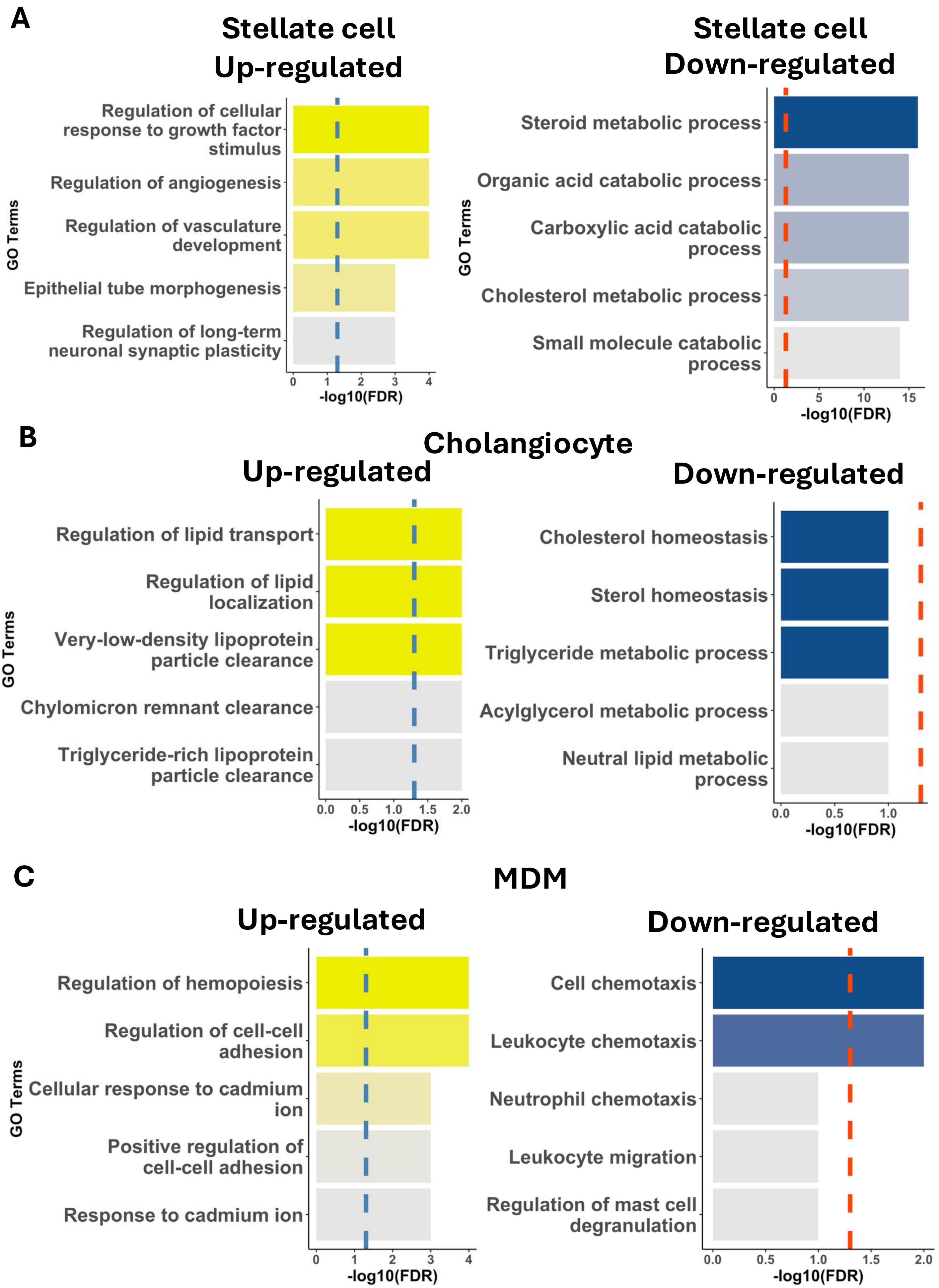
A
Stellate cell
Up-regulated
Regulation of cellular response to growth factor stimulus
Regulation of angiogenesis
Regulation of vasculature development
Epithelial tube morphogenesis
Regulation of long-term neuronal synaptic plasticity
GO Terms
0
1
2
3
4
-log10(FDR)
Stellate cell
Down-regulated
Steroid metabolic process
Organic acid catabolic process
Carboxylic acid catabolic process
Cholesterol metabolic process
Small molecule catabolic process
GO Terms
0
5
10
15
-log10(FDR)
B
Cholangiocyte
Up-regulated
Regulation of lipid transport
Regulation of lipid localization
Very-low-density lipoprotein particle clearance
Chylomicron remnant clearance
Triglyceride-rich lipoprotein particle clearance
GO Terms
0.0
0.5
1.0
1.5
2.0
-log10(FDR)
Down-regulated
Cholesterol homeostasis
Sterol homeostasis
Triglyceride metabolic process
Acylglycerol metabolic process
Neutral lipid metabolic process
0.0
0.5
1.0
-log10(FDR)
C
MDM
Up-regulated
Regulation of hemopoiesis
Regulation of cell-cell adhesion
Cellular response to cadmium ion
Positive regulation of cell-cell adhesion
Response to cadmium ion
GO Terms
0
1
2
3
4
-log10(FDR)
Down-regulated
Cell chemotaxis
Leukocyte chemotaxis
Neutrophil chemotaxis
Leukocyte migration
Regulation of mast cell degranulation
0.0
0.5
1.0
1.5
2.0
-log10(FDR)

### Fig. S2

**Figure S2**. Top 5 down-regulated gene ontology enrichment results in endothelial cell stellate cell (**A**), cholangiocyte (**B**), and MDM (**C**) from perinatal exposure to PCBs.  Dotted lines represent -log10 FDR-adjusted p-value at 0.05.

Fig. S3

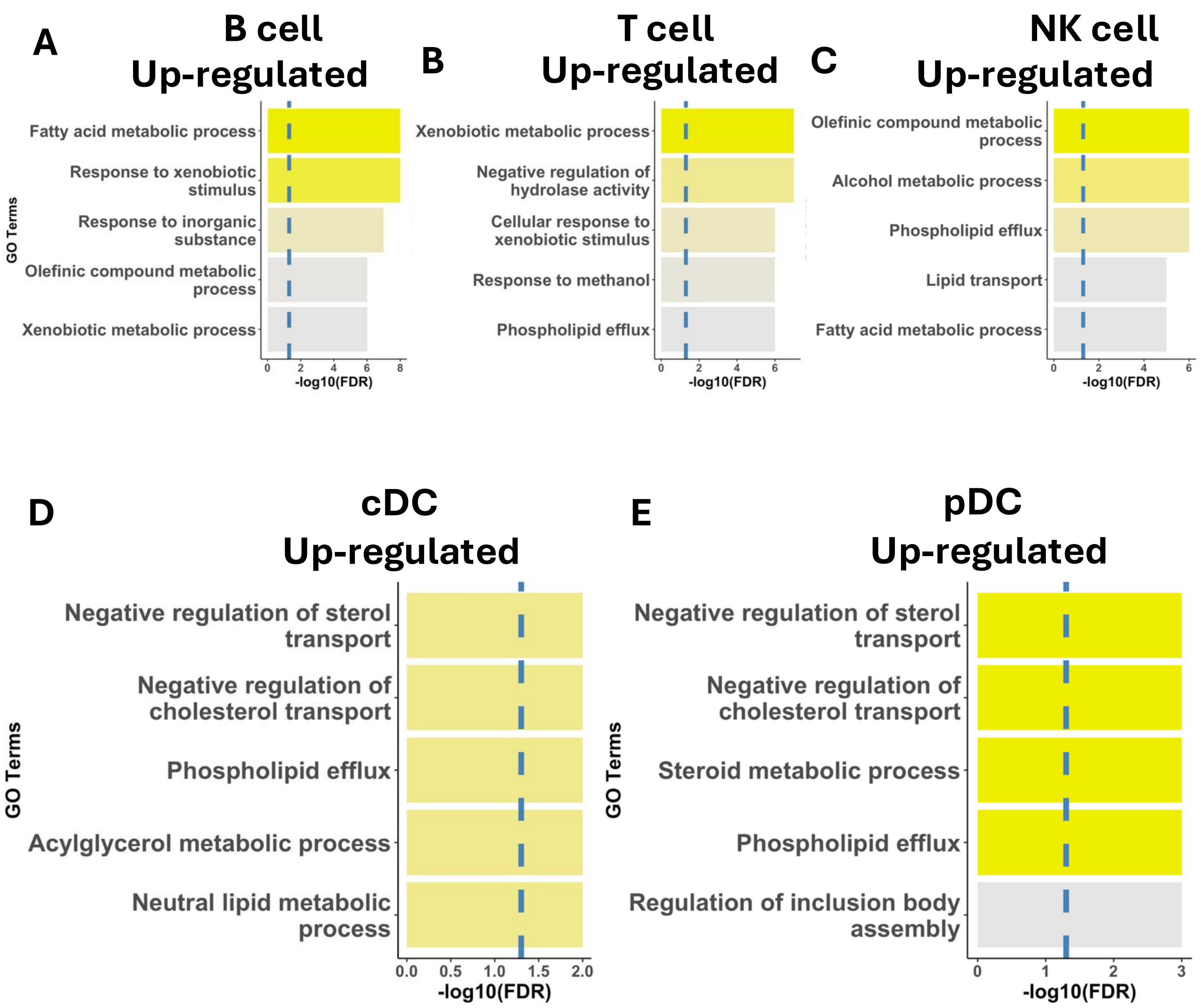


**Figure S3**. Top 5 up-regulated gene ontology enrichment results in B cell (**A**), T cell (**B**), NK cell (**C**), cDC (**D**), and pDC (**E**) from perinatal exposure to PCBs. Dotted lines represent -log10 FDR-adjusted p-value at 0.05.

Fig. S4

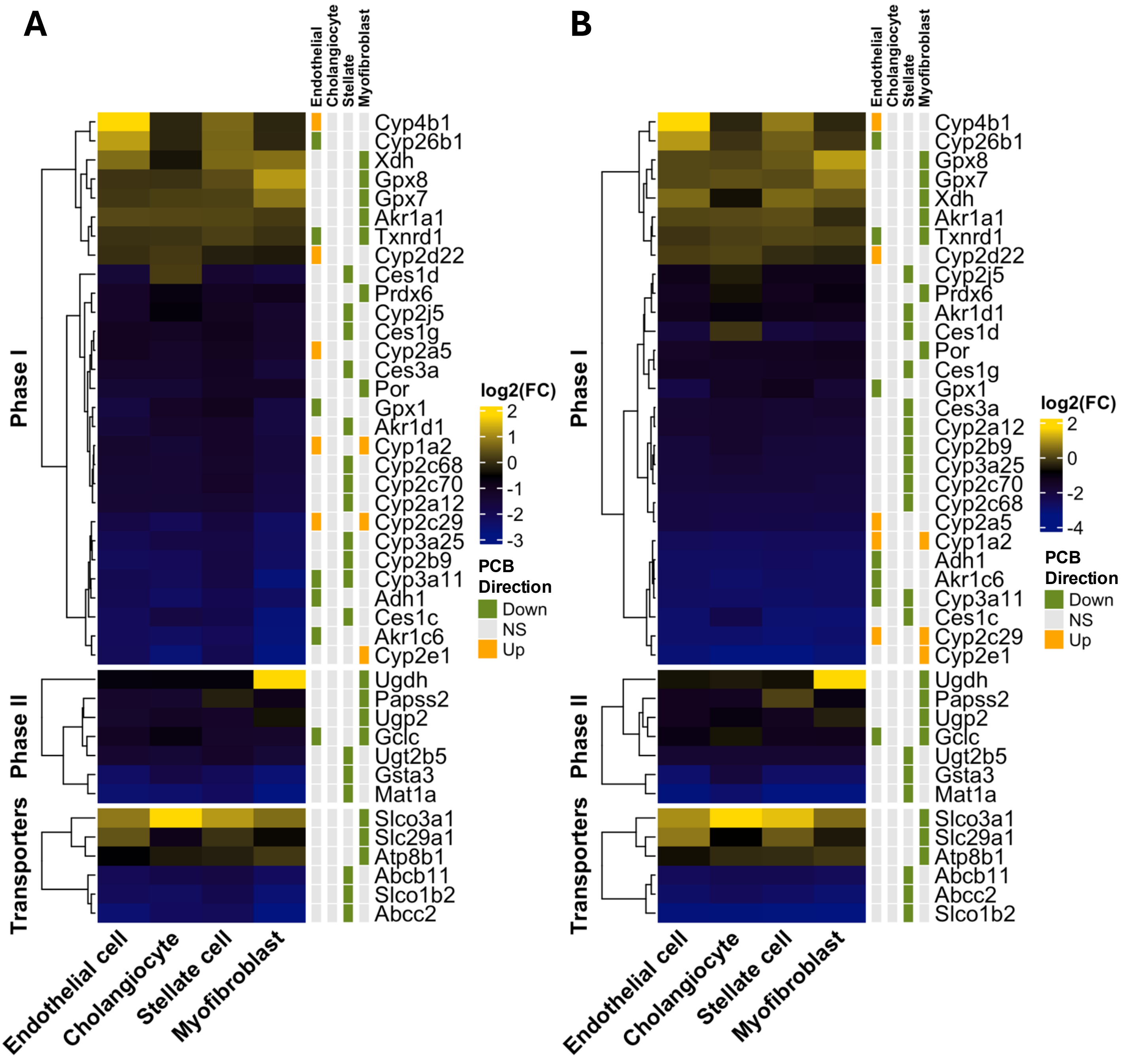


**Figure S4.** Log2 fold change values in endothelial cells, cholangiocytes, stellate cells, and myofibroblasts for persistently dysregulated drug processing genes relative to hepatocytes in the vehicle (**A**) and PCB-exposed group (**B**). Yellow and blue in the heatmap represent the log2 fold change of liver genes of the PCB exposed adults in non-parenchymal cells with respect to hepatocytes. "PCB Direction" indicates whether a gene is up- or down-regulated from perinatal exposure to PCB (Bonferroni-adjusted p-value < 0.05).

**Fig. S5**

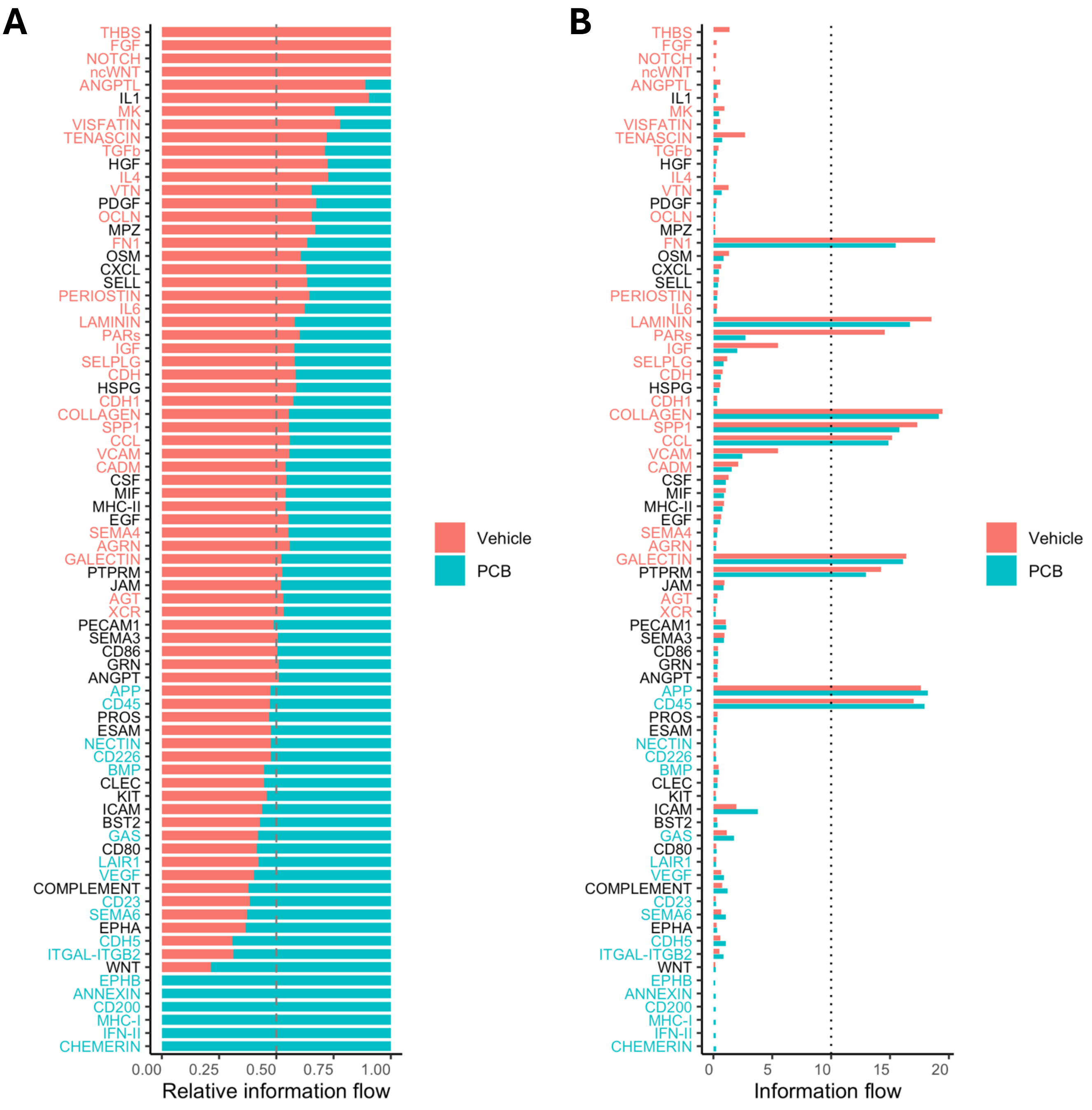


**Figure S5.** Predicted intercellular communication results. **A**. Relative change of predicted signaling pathways in vehicle (red) or PCBs (blue). The predicted signaling patterns are shown in the y axis. Red or blue indicates significantly enriched predicted signaling pathways by vehicle or PCBs, respectively. **B**. Total log probability of intercellular signaling signaling (x axis) by each signaling pathway (y axis). Dotted line shows threshold value at 10. Red or blue indicates significantly enriched predicted signaling pathways by vehicle or BDE-99, respectively.

**Fig. S6**

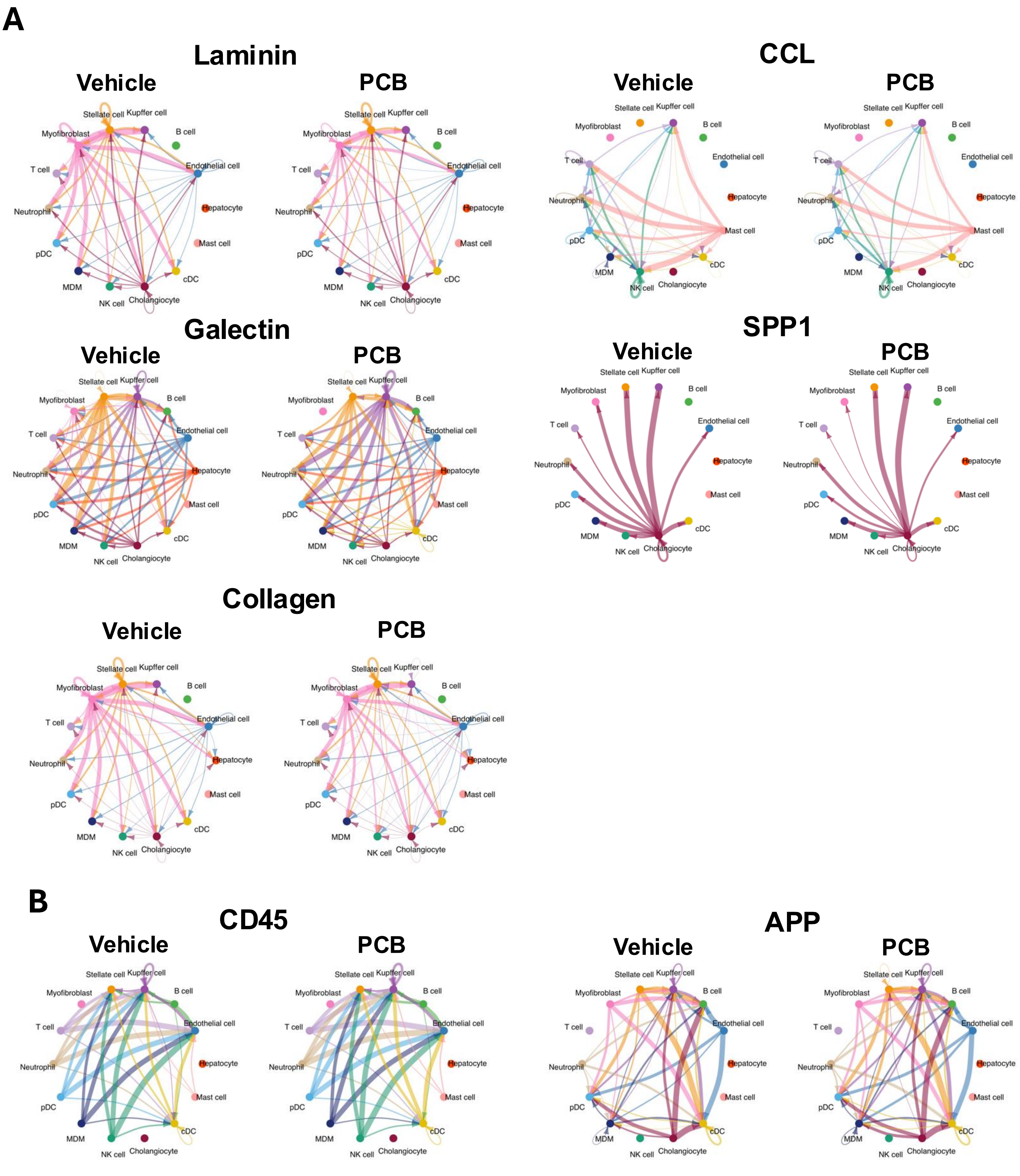


**Figure S6**. Visualization of the cell-cell communications of the signaling pathways that are predicted to be decreased (**A**) or increased (**B**) in PCB exposed pups. Each cell type contains a unique color, and the matched colors represent signal communication direction from one cell type to another. The thickness of the arrows represents the probability of communication.
